# A microbiota-dependent gut bile acid axis links germline mutant p53 to distal tumour growth

**DOI:** 10.64898/2026.10.05.753853

**Authors:** Noel Wei Yang Ong, Camilla Giovino, Paula Rosanna Quaglietta, Yoonseo Park, Raya Traubici, Nicholas William Fischer, David Malkin

**Affiliations:** Department of Medical Biophysics, University of Toronto; Toronto, Ontario, Canada; Program in Genetics and Genome Biology, The Hospital for Sick Children; Toronto, Ontario, Canada; Department of Biochemistry, Seoul National University; Seoul, Republic of Korea; Institute for Management & Innovation, University of Toronto Mississauga; Mississauga, Ontario, Canada; Division of Haematology/Oncology, The Hospital for Sick Children; Toronto, Ontario, Canada; Department of Paediatrics, University of Toronto; Toronto, Ontario, Canada

**Author notes:** Corresponding author: David Malkin.

## Abstract

The gut microbiome can influence cancer development, but its contribution to cancer predisposition in Li-Fraumeni Syndrome (LFS) remains poorly understood. Here, we investigated whether germline mutant p53 alters the intestinal environment using *Trp53*^R172H/+^ mice. Tumour-naïve mutant mice exhibited increased intestinal permeability and a pro-inflammatory colonic phenotype, including elevated inflammatory cytokines and enhanced NF-κB signalling. Untargeted metabolomic profiling revealed a pronounced alteration in bile acid metabolism, with targeted analysis identifying increases in multiple primary and microbiota-derived secondary bile acids. Interestingly, these metabolic changes occurred without detectable shifts in overall microbial community composition, suggesting that host-microbial metabolic interactions may be altered independently of broad changes in microbiota composition. Several faecal bile acids were positively associated with distal tumour burden, with the microbiota-derived secondary bile acid deoxycholic acid (DCA) showing the strongest relationship. Microbiota depletion reduced distal tumour growth in mutant mice and attenuated intestinal barrier dysfunction, colonic inflammation and the altered faecal bile acid profile, including DCA. Conversely, transfer of cell-free faecal filtrate from mutant donors to WT recipients induced colonic inflammation and enhanced distal tumour growth. Together, these findings uncover a previously unrecognized mechanism by which germline mutant p53 reshapes the intestinal metabolic and inflammatory landscape, creating a microbiota-dependent environment that fuels tumour growth at distant sites and may contribute to cancer predisposition in LFS.

## Introduction

Li-Fraumeni Syndrome (LFS) is a highly penetrant cancer-predisposition disorder caused by pathogenic germline *TP53* variants^1–4^. Although this susceptibility is primarily attributed to impaired tumour suppression within emerging malignant cells, germline *TP53* variants are present throughout the host and may also alter non-malignant tissues in ways that favour tumour development. Mutant p53 (mutp53) can influence inflammatory signalling, metabolism and the immune microenvironment^5–8^, but whether germline mutp53 creates a tumour-permissive host environment remains poorly understood.

The intestine is a major interface between host genetics, immunity, and the microbiota^9^. Wild-type p53 (wtp53) can restrain NF-κΒ signalling^10,11^, whereas mutp53 has been shown to prolong NF-κΒ activation and promote chronic inflammation in the colon^5^. Mucosal inflammatory signalling is closely linked to intestinal barrier integrity, and barrier dysfunction can increase host exposure to microbial products and metabolites, further amplifying inflammation^12–14^. Conversely, microbiota-derived metabolites can also alter mutp53 activity in the intestine and shift its function toward an oncogenic state^15^. Beyond the gut, the intestinal microbiota can influence tumours at anatomically distant sites, including through host genetic control of the intestinal environment ^16–18^. Despite these established interactions, whether germline mutp53 drives intestinal metabolic and inflammatory changes that contribute to tumour development beyond the gut remains unknown.

Bile acids are potential mediators of host-microbial interactions, linking host cholesterol metabolism with microbial metabolic activity. Primary bile acids synthesized from cholesterol are modified by intestinal bacteria to generate secondary bile acids^19–21^, several of which have pro-inflammatory and pro-tumourigenic properties^22–24^. Bile acids act on host cells largely through dedicated receptors, chiefly the nuclear receptor FXR and the membrane receptor TGR5, both of which shape intestinal barrier function and inflammatory tone ^25,26^. p53 has emerged as a regulator of mevalonate and cholesterol metabolism, with wtp53 repressing and mutp53 enhancing pathway activity^6,27^. Thus, germline *TP53* mutations may influence both host metabolic pathways and microbiota-dependent bile acid metabolism, potentially generating a pro-inflammatory intestinal environment that favours tumour development at distant sites.

Here, using *Trp53*^R172H/+^ mice carrying the murine equivalent of the human *TP53* R175H hotspot mutation^28^, we examined intestinal barrier function, mucosal inflammation, microbial composition, and metabolic output. We also assessed how these intestinal features relate to distal tumour growth following subcutaneous tumour challenge. We used antibiotic-mediated microbiota depletion to determine the contribution of the microbiota to the altered intestinal state and enhanced tumour growth, and cell-free faecal filtrate transfer to test whether transferable faecal components were sufficient to reproduce these phenotypes ^14^. Our findings reveal that germline mutp53 establishes a microbiota-dependent inflammatory and metabolic intestinal environment that promotes tumour growth at distant sites, uncovering a previously unrecognized dimension of cancer predisposition in LFS.

## Results

### Mutant p53 compromises the intestinal barrier function and activates pro-inflammatory NF-κB signalling in the colon

Given the gastrointestinal tract’s role as a major interface between environmental cues and host immunity, and evidence that the microbial environment can modulate mutp53 function^15^, we examined whether germline mutp53 was associated with altered intestinal barrier function and mucosal inflammation. We first assessed intestinal barrier integrity using an *in vivo* permeability assay. Oral gavage of 4 kDa FITC-dextran followed by plasma fluorescence quantification revealed significantly higher circulating FITC-dextran concentrations in mutant mice than in WT littermates (p = 0.0249; Fig. 1A). Despite this functional barrier defect, crypt length was unchanged between genotypes (p = 0.2254; Fig. 1B, C), and overall tissue architecture appeared grossly normal by haematoxylin and eosin staining. These findings indicate that increased intestinal permeability occurs in the absence of overt histological abnormalities.

**Fig. 1.**
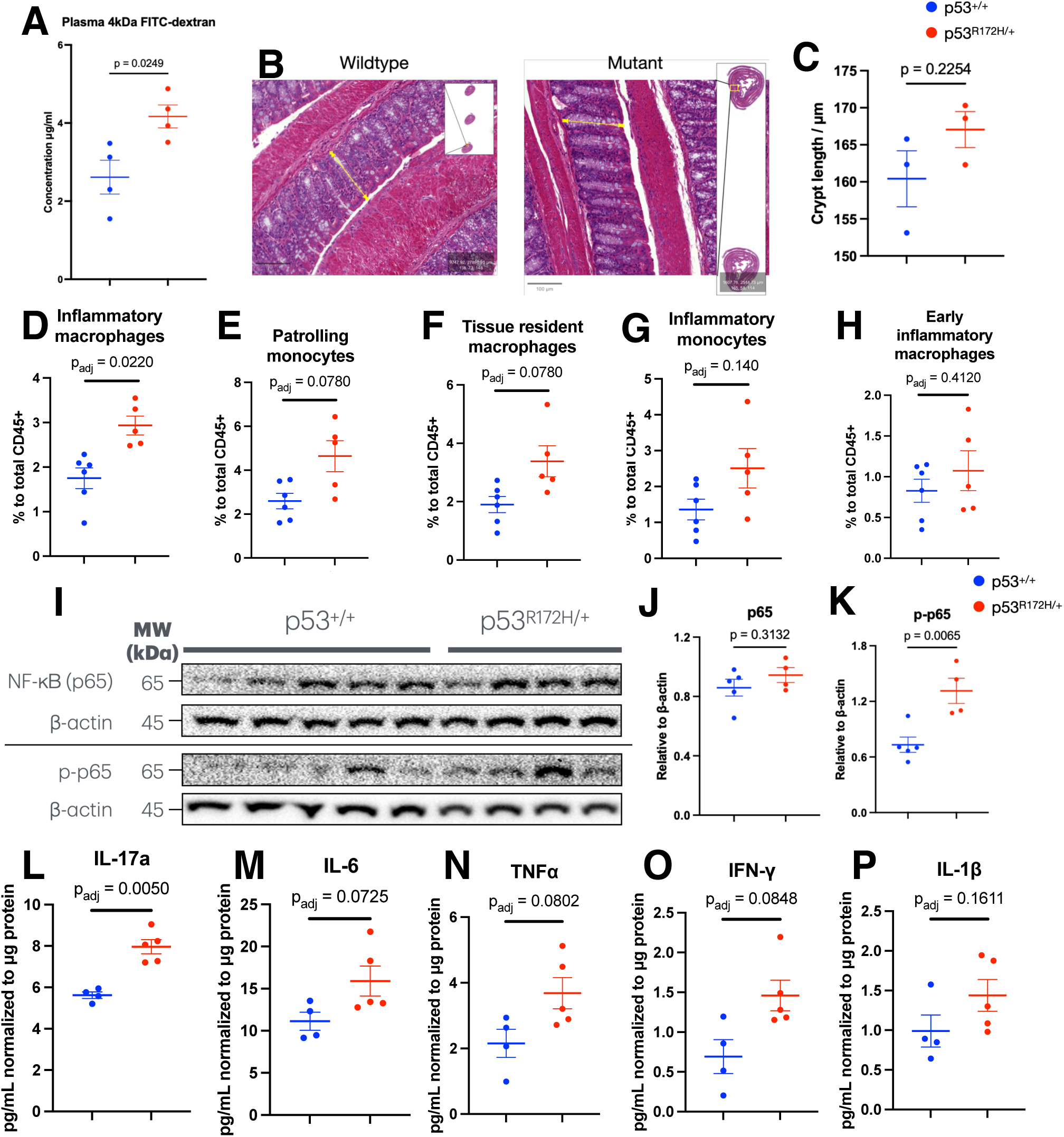
Mutant p53 compromises intestinal barrier integrity and activates pro-inflammatory NF-κB signalling in the colon. (A) Plasma concentration of orally gavaged 4-kDa FITC-dextran in *Trp53*^+/+^ (WT) and *Trp53*^R172H/+^ (mutant) mice. (B) Representative haematoxylin and eosin-stained sections of the distal colonic mucosa from WT and mutant mice; yellow lines indicate the crypt-length measurements quantified in C. (C) Colonic crypt length in WT and mutant mice. (D–H) Flow cytometric quantification of colonic lamina propria myeloid subsets (gated on live CD45^+^CD19^-^ cells), shown as percentage of total CD45^+^ cells: (D) inflammatory macrophages (Ly6C^hi^F4/80^+^CD11b^+^CD11c^+^); (E) patrolling monocytes (Ly6C^lo^F4/80^-^CD11b^+^CD11c^-^); (F) tissue resident macrophages (Ly6C^lo^F4/80^+^CD11b^+^CD11c^+^); (G) inflammatory monocytes (Ly6C^hi^F4/80^-^CD11b^+^CD11c^-^); (H) early inflammatory macrophages (Ly6C^hi^F4/80^+^CD11b^+^CD11c^-^). (I) Representative immunoblots of colonic lysates probed for total NF-κB (p65) and phosphorylated p65 (p-p65), with β-actin as a loading control; molecular weights (kDa) are indicated. (J, K) Densitometric quantification of total p65 (J) and p-p65 (K) in I, normalized to β-actin. (L-P) Colonic tissue cytokine concentrations (normalized to μg total protein), measured by ELISA: (L) IL-17a; (M) IL-6; (N) TNFα; (O) IFN-γ; (P) IL-1β. Data are mean ± SEM; each symbol represents one biologically independent mouse (n = 4–6 mice per genotype; blue, *Trp53*^+/+^; red, *Trp53*^R172H/+^). Group differences were assessed by two-tailed Welch’s t-test. For D-H and L-P, p values were adjusted for multiple comparisons across the five myeloid populations and the five cytokines, respectively, using the Benjamini-Hochberg procedure at a false discovery rate of 5%. Flow cytometry samples were acquired once per mouse; ELISA samples were assayed in duplicate wells and averaged before analysis.

Consistent with an inflammatory intestinal phenotype, flow cytometric analysis of the colonic lamina propria point in the same direction. Within the live CD45^+^CD19^-^ compartment, inflammatory macrophages were higher in mutant mice than in WT littermates and were the only subset for which this difference survived correction for multiple comparisons (p_adj_ = 0.0220; Fig. 1D). Patrolling monocytes (p_adj_ = 0.0780; Fig. 1E), tissue resident macrophages (p_adj_ = 0.0780; Fig. 1F), and inflammatory monocytes (p_adj_ = 0.1400; Fig. 1G) were each elevated in mutant mice without reaching significance, and early inflammatory macrophages were present at comparable frequencies in the two genotypes (p_adj_ = 0.4120; Fig. 1H). The shift was therefore graded rather than uniform: it was largest in the most differentiated population sampled, absent in the earliest precursor, and intermediate in between. With five to six mice per genotype this experiment cannot resolve the smaller differences, and we read the subthreshold trends as suggestive only. The gradient is nonetheless more consistent with directed recruitment and local differentiation of monocyte-derived cells than with a generalized myeloid influx.

Given previous evidence that mutp53 can prolong NF-κΒ activation in the colon, we next examined NF-κΒ pathway activation in colonic tissue ^5^. Immunoblot analysis showed that total NF-κΒ (p65) levels remained unchanged (p = 0.3132), whereas phosphorylated p65 was significantly increased in mutant mice (p = 0.0065; Fig. 1I-K), indicating heightened pathway activity. Cytokine measurements in colonic tissue lysates were similarly selective. IL-17a was significantly elevated in mutant mice (p_adj_ = 0.0050; Fig. 1L), whereas IL-6 (p_adj_ = 0.0725; Fig. 1M), TNF-α (p_adj_ = 0.0802, Fig. 1N), IFN-γ (p_adj_ = 0.0848; Fig. 1O), and IL-1β (p_adj_ = 0.1611; Fig. 1P) were each higher in mutant tissue but did not reach significance after correction across the panel. That all five analytes moved in the same direction is less easily attributed to chance than any one comparison suggests, although the present cohort is too small to test this formality. Mutant mice thus displayed an intestinal phenotype of barrier dysfunction, a shifted mucosal myeloid compartment and heightened NF-κΒ signalling, with IL-17a the most clearly affected cytokine.

### Mutant p53 is associated with altered intestinal bile acid metabolism

To investigate whether the inflammatory intestinal phenotype in mutp53 mice was accompanied by changes in the gut metabolic environment, we performed untargeted metabolomic profiling of faecal samples. Bile acid metabolism emerged as the top-ranked enriched pathway in mutant mice (Fig. 2A). This finding was notable given the established role of p53 in mevalonate and cholesterol metabolism and the pro-inflammatory and pro-tumourigenic properties of several bile acid species ^6,22,23,27^.

**Fig. 2.**
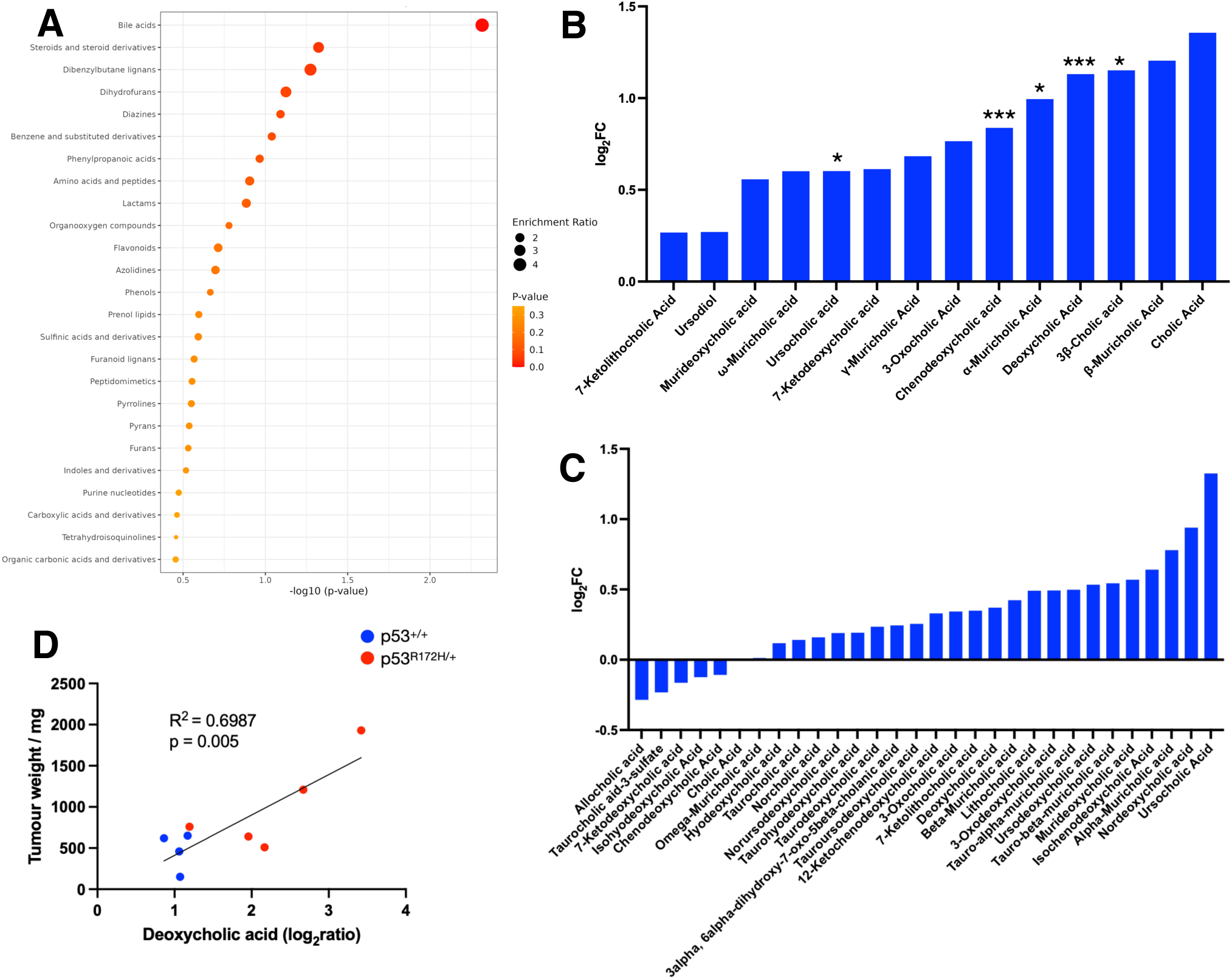
Mutant p53 drives accumulation of pro-tumourigenic secondary bile acids in the gut. (A) Bubble plot of metabolite-class enrichment from untargeted faecal metabolomic profiling of WT and mutant mice, generated using MetaboAnalyst ^43^. X-axis, −log_10_(*p* value); bubble size, enrichment ratio; colour, *p* value. Bile acid metabolism was the most significantly enriched pathway. (B) Log_2_ fold-change (mutant/WT) of individual faecal bile acid species. (C) Log_2_ fold-change (mutant/WT) of individual plasma bile acid species; no individual species reached statistical significance. (D) Correlation between faecal deoxycholic acid abundance (log_2_ ratio, mutant/WT) and subcutaneous tumour weight following MC38 colon adenocarcinoma challenge; linear regression with 95% confidence interval. n = 4–5 faecal and plasma samples per genotype. In B, asterisks denote species with statistically significant differences (one asterisk, *p*_adj_ < 0.05; two asterisks, *p*_adj_ < 0.01; three asterisks, *p*_adj_ < 0.001), corrected for multiple comparisons; exact *R*² and *P* values for D are shown on the graph.

Targeted analysis confirmed significant elevations across multiple primary and secondary bile acids in the faeces of mutant mice relative to WT mice (Fig. 2B and Supplementary Fig. S1). Several of the elevated species were microbiota-modified bile acids, including deoxycholic acid (DCA), a secondary bile acid with established tumour-promoting effects^23,29^. Plasma bile acid profiles also showed an overall increase in mutp53 mice, with many secondary bile acids among the positively shifted species (Fig. 2C, and Supplementary Fig. S2). Although individual plasma bile acids did not reach significance, this pattern was consistent with the more pronounced changes detected within the intestinal compartment.

To determine whether this altered bile acid profile was associated with tumour growth, we correlated faecal bile acid levels with tumour weight following MC38 tumour challenge. Multiple faecal bile acids showed positive correlations with the tumour burden, with DCA showing the strongest positive correlation (R^2^ = 0.6987, p = 0.005; Fig. 2D). These findings identify altered bile acid metabolism, including increases in microbiota-modified species, as a prominent feature of the mutp53 intestinal environment.

### Gut microbiota composition remains stable despite functional metabolic changes

Given the marked alterations in bile acids, we next examined whether mutp53 was associated with shifts in the gut microbial population. 16S rRNA sequencing revealed no significant differences in α-diversity metrics (Shannon, p = 0.328; Simpson, p = 0.313; Chao-1, p = 0.515; observed richness indices, p = 0.524) between WT and mutant mice (Fig. 3A). Principal coordinates analysis of Bray-Curtis dissimilarity confirmed that community level β-diversity remained comparable between genotypes (PERMANOVA, R^2^ = 0.007, p = 0.723), with no distinct clustering observed (Fig. 3B). Taxonomic relative abundance profiles and genus-level analyses further demonstrated a stable microbial landscape regardless of p53 status (Fig. 3C, and Supplementary Fig. S3, S4). The presence of marked bile acid alterations in the absence of detectable changes in microbial community composition suggests that mutp53 may alter the metabolic activity of the gut microbiota without broadly reshaping the microbial community.

**Fig. 3.**
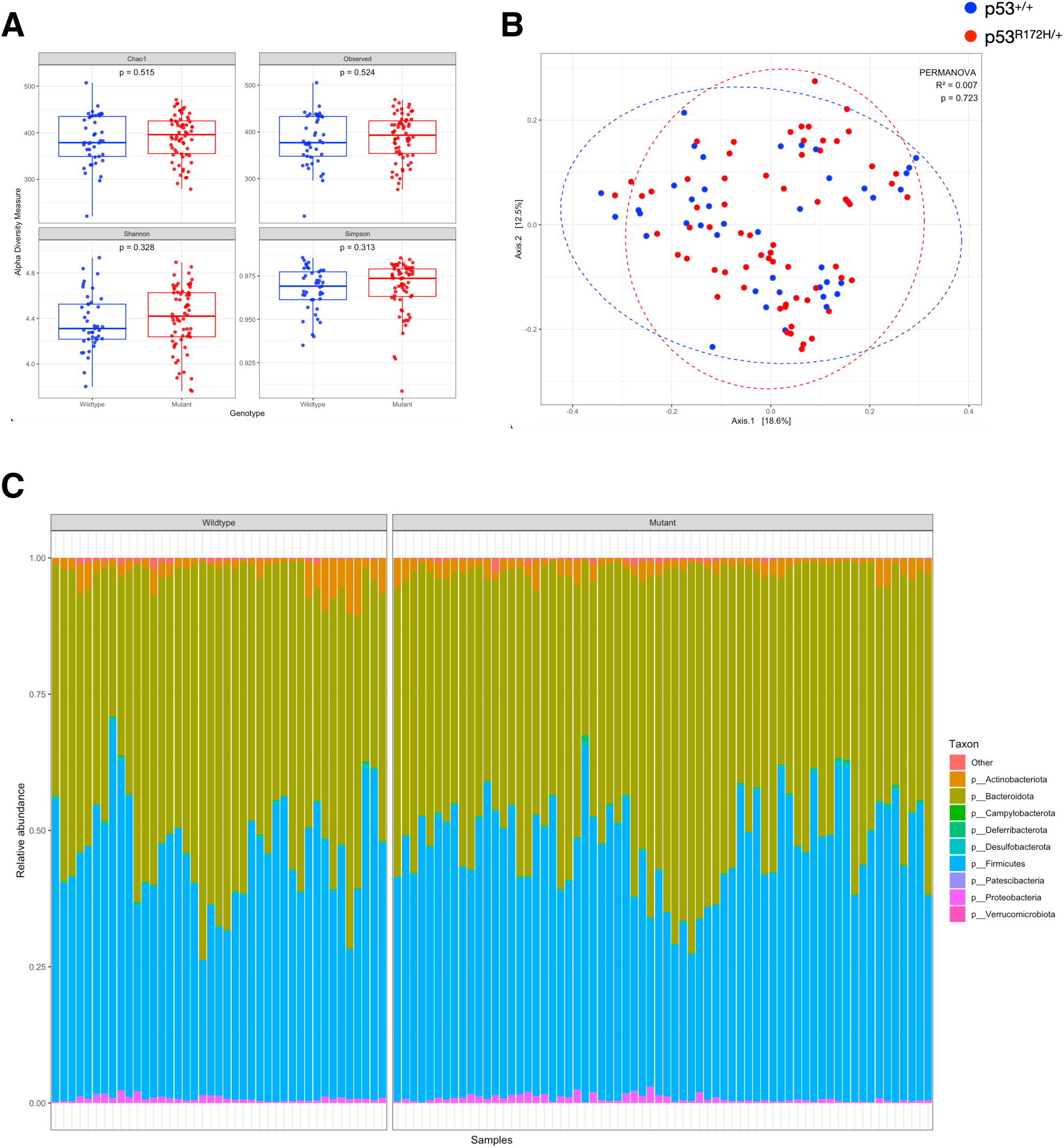
Gut microbiota composition is not significantly altered by mutant p53 despite functional metabolic changes. (A) α-diversity of faecal microbiota in WT and mutant mice (Chao1, observed richness, Shannon and Simpson indices), determined by 16S rRNA gene sequencing (V4 hypervariable region, Illumina MiSeq); none of the indices differed significantly (two-tailed Wilcoxon rank-sum test). (B) Principal coordinates analysis (PCoA) of Bray–Curtis dissimilarity comparing faecal microbial community composition (β-diversity) between WT and mutant mice; dashed ellipses indicate 95% confidence intervals. Community-level differences were assessed by PERMANOVA. (C) Relative abundance of bacterial phyla in individual faecal samples from WT and mutant mice. n = 41–66 mice per genotype.

### Microbiota depletion suppresses distal non-intestinal tumour growth and abrogates colonic inflammation

To test whether the gut microbiota contributes causally to the mutp53 phenotype, we administered a broad-spectrum antibiotic cocktail to mutant and WT mice before and during subcutaneous MC38 challenge. Microbiota depletion markedly reduced subcutaneous tumour mass in mutant mice compared with vehicle-treated mutant controls (p_adj_ = 0.0010; Fig. 4A). In contrast, antibiotic treatment had no significant effect on tumour growth in WT littermates (p_adj_ = 0.9967), consistent with a tumour-promoting contribution of the microbiota that is specific to the mutant p53 background.

**Fig. 4.**
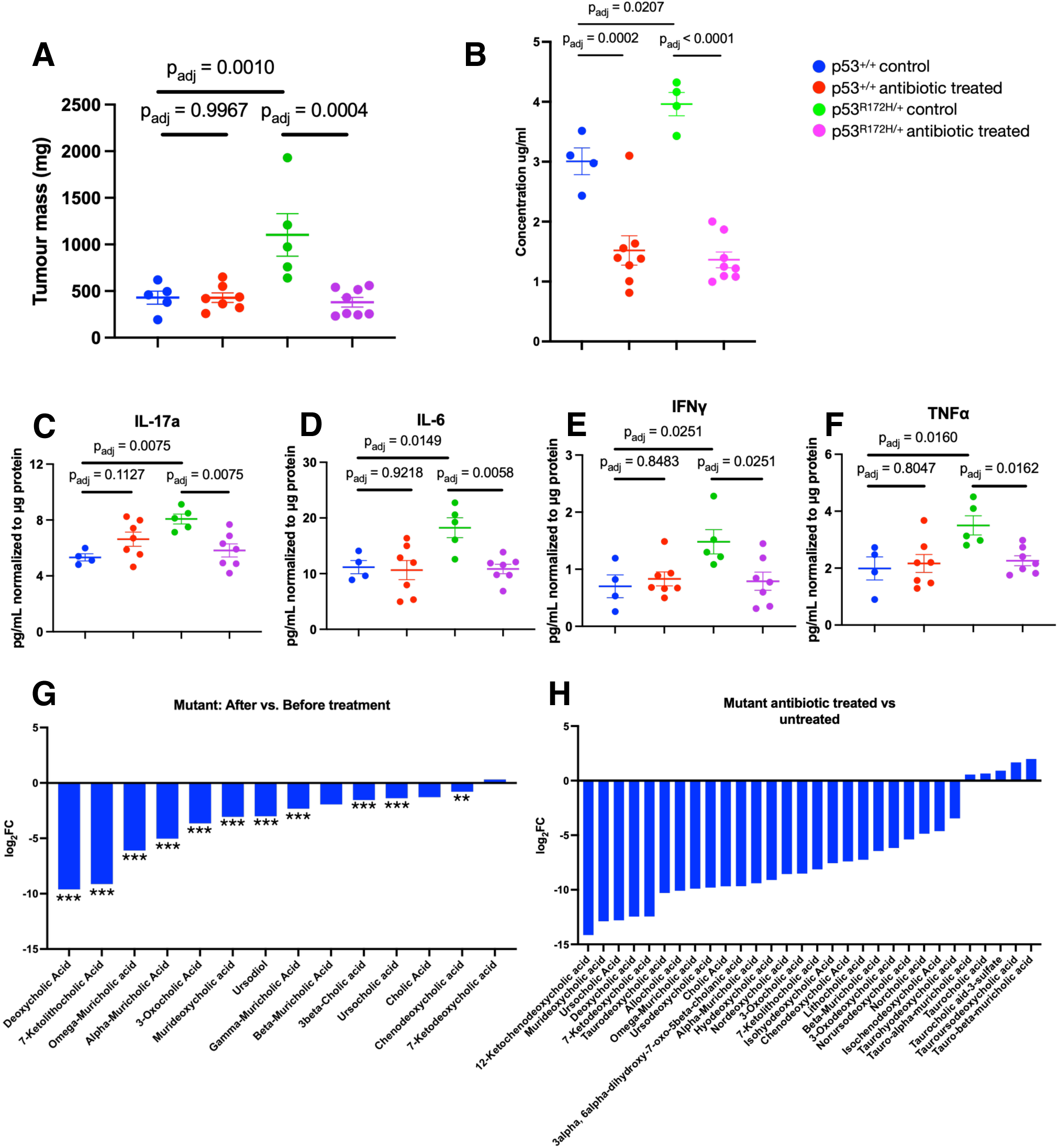
Microbiota depletion suppresses distal tumourigenesis and reverses colonic barrier dysfunction and inflammation in mutant mice. (A) Subcutaneous tumour mass in *Trp53*^+/+^ control (blue), *Trp53*^+/+^ antibiotic-treated (red), *Trp53*^R172H/+^ control (green) and *Trp53*^R172H/+^ antibiotic-treated (magenta) mice following oral administration of a broad-spectrum antibiotic cocktail (composition in Methods). (B) Plasma 4-kDa FITC-dextran concentration across the same four groups. (C-F) Colonic tissue cytokine concentrations (normalized to μg total protein) across the same four groups: (C) IL-17a; (D) IL-6; (E) IFN-γ; (F) TNFα. (G) Log_2_ fold-change of individual bile acid species in mutant mice after versus before antibiotic treatment. (H) Log_2_ fold-change of individual plasma bile acid species in antibiotic-treated versus untreated mutant mice; no individual species reached statistical significance. Data are mean ± SEM; each symbol represents one biologically independent mouse (n = 4–8 mice per genotype). Adjusted *p* values for (A-F) were determined by one-way ANOVA, corrected with Benjamini–Hochberg correction at a false discovery rate of 5%; exact values are reported in the Results. In (G), asterisks denote species with statistically significant differences (one asterisk, *p*_adj_ < 0.05; two asterisks, *p*_adj_ < 0.01; three asterisks, *p*_adj_ < 0.001), corrected for multiple comparisons.

Barrier function and colonic inflammation responded differently to depletion, and the difference is informative. Antibiotic treatment reduced plasma FITC-dextran in both genotypes (WT, p_adj_ = 0.0002; mutant, p_adj_ < 0.0001), on top of the expected baseline difference between mutant and WT controls (p_adj_ = 0.0207; Fig. 4B). Microbiota depletion also attenuated the inflammatory colonic phenotype, significantly reducing IL-17a (p_adj_ = 0.0075), IL-6 (p_adj_ = 0.0149), IFN-γ (p_adj_ = 0.0251), and TNF-α (p_adj_ = 0.0160) in mutant mice (Fig. 4C-F). These cytokines were not significantly altered by antibiotic treatment in WT mice, supporting a microbiota-dependent inflammatory phenotype associated with mutant p53.

Untargeted metabolomic analysis showed that antibiotic treatment substantially decreased the abundance of multiple faecal bile acids in mutant mice (Fig. 4G). Notably, DCA was significantly reduced following antibiotic treatment, alongside several other bile acid species (Supplementary Fig. S5). Plasma bile acids showed a similar downward trend, although these changes did not reach statistical significance (Fig. 4H and Supplementary Fig. S6). Together, microbiota depletion attenuated the mutp53-associated intestinal metabolic and inflammatory phenotype in parallel with reduced distal tumour growth.

### Transfer of mutant p53-associated faecal milieu promotes colonic inflammation and distal tumour growth

Having established a contribution of the microbiota to the mutp53 phenotype, we next examined whether cell-free components of the altered intestinal environment were sufficient to reproduce the enhanced tumour growth and colonic inflammatory phenotype in WT mice. We therefore performed faecal filtrate transplant (FFT) from mutant donors to WT recipients. This approach transfers soluble and filterable faecal components while excluding live bacterial transfer.

WT recipients of mutp53-associated faecal filtrate developed significantly larger subcutaneous tumours than vehicle-treated WT controls (p_adj_ = 0.0282; Fig. 5A). This increase in distal tumour growth was accompanied by a broad inflammatory response in the colon. Mutant faecal filtrate significantly increased colonic IFN-γ (p_adj_ < 0.0001), IL-1β (p_adj_ = 0.0021), IL-17a (p_adj_ = 0.0250), TNF-α (p_adj_ = 0.0367), and IL-6 (p_adj_ = 0.0428) (Fig. 5B-F), recapitulating the broad colonic cytokine elevation observed in untreated mutant mice.

**Fig. 5.**
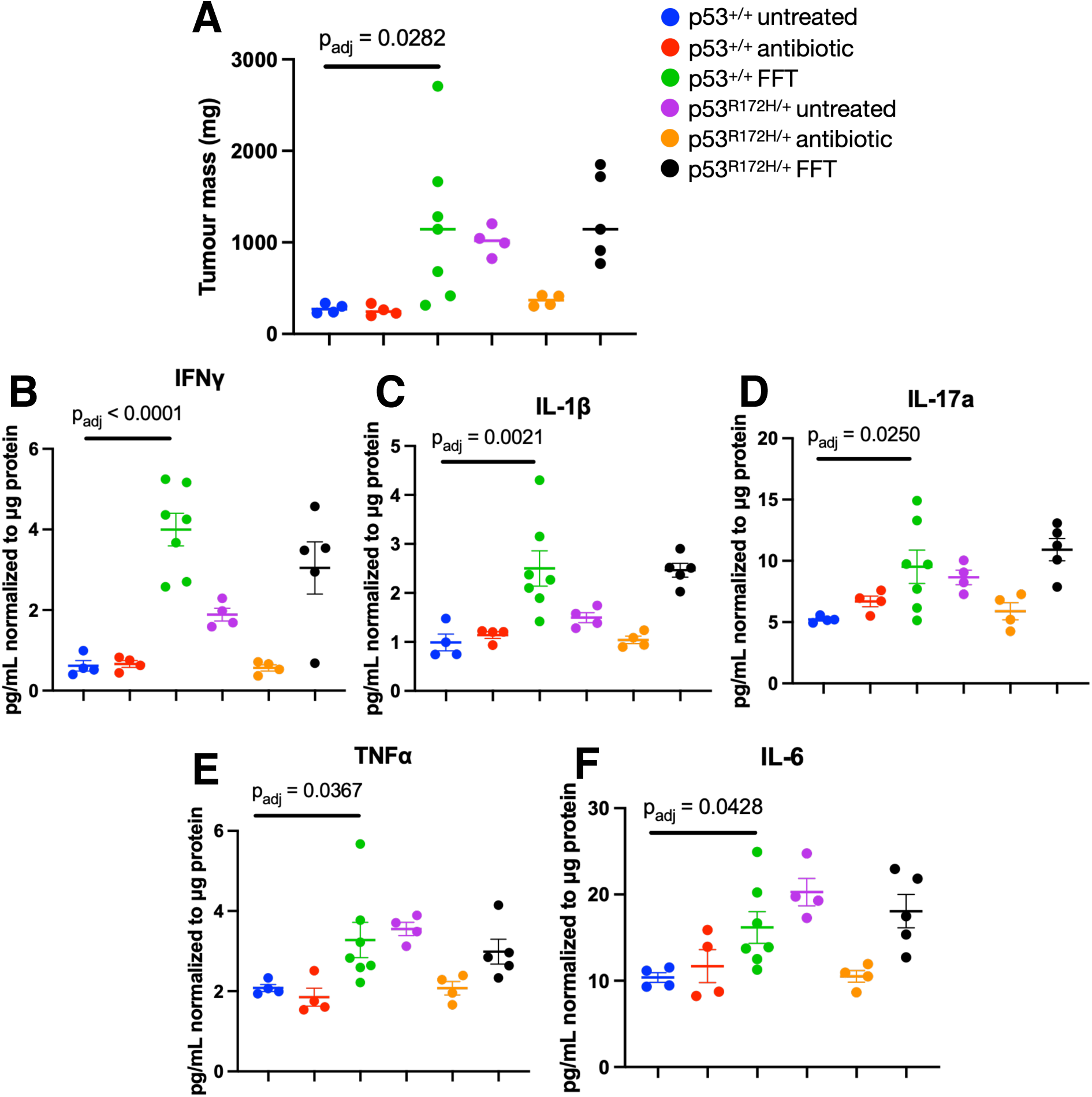
Transfer of mutp53-associated faecal filtrate is sufficient to recapitulate colonic inflammation and accelerate distal tumourigenesis in wild-type mice. (A) Subcutaneous tumour mass in WT and mutant recipients that were untreated, antibiotic-depleted or received faecal filtrate transplantation (FFT) from mutant donor mice. (B-F) Colonic tissue cytokine concentrations (normalized to μg total protein) across the same treatment groups: (B) IFN-γ; (C) IL-1β; (D) IL-17a; (E) TNFα; (F) IL-6. Data are mean ± SEM; each symbol represents one biologically independent mouse (n = 4–5 mice per group). Adjusted *p* values were determined by one-way ANOVA with Benjamini–Hochberg correction; the comparison of interest throughout is between the WT untreated and WT FFT groups. Exact values are reported in the Results.

Together, these findings demonstrate that cell-free components of the mutp53-associated faecal milieu are sufficient to promote colonic inflammation and enhance distal tumour growth in WT mice, supporting the presence of transferable mediators within the altered intestinal environment.

## Discussion

Our findings identify intestinal barrier dysfunction and altered bile acid metabolism as previously unrecognized features of the tumour-permissive host environment associated with germline mutant p53. Mutp53 mice exhibited increased intestinal permeability, altered colonic myeloid populations, enhanced NF-κB signalling and elevated inflammatory cytokines, together with increases in faecal bile acids despite no detectable differences in overall microbial community composition. Microbiota depletion attenuated these intestinal abnormalities and reduced distal tumour growth, whereas transfer of mutp53-associated faecal filtrate promoted colonic inflammation and tumour growth in WT recipients. Together, these findings extend the consequences of germline *TP53* mutations beyond tumour-cell-intrinsic defects and support a model in which mutp53 establishes an altered intestinal metabolic and inflammatory state that is shaped by the microbiota and capable of promoting tumour growth beyond the gut.

The increased NF-κB activity observed in the mutp53 colon is consistent with previous evidence that mutant p53 can prolong NF-κB signalling and promote intestinal inflammation^5^. Here, increased p65 phosphorylation occurred alongside elevated IL-17a, enrichment of mucosal myeloid populations, and increased intestinal permeability despite the absence of overt histological abnormalities. This combination of barrier impairment and IL-17-associated inflammation resembles the axis through which microbial products drive tumour growth in other intestinal models ^30^. Microbiota depletion reduced intestinal permeability in both genotypes, whereas significant reductions in multiple colonic cytokines were observed in mutp53 mice but not WT mice, indicating that the microbial environment contributes to barrier function and to the inflammatory state associated with mutp53.

Untargeted metabolomic profiling identified bile acid metabolism as the most prominently altered metabolic pathway, with targeted analysis demonstrating elevations across multiple primary and secondary bile acids. This finding is particularly relevant because cholesterol is the precursor for primary bile acid synthesis and p53 regulates mevalonate and cholesterol metabolism^6,27^. WT p53 has also been shown to directly regulate bile acid homeostasis through activation and stabilization of Small Heterodimer Partner (SHP), an inhibitor of bile acid synthesis, providing an additional link between p53 status and regulation of the bile acid pool^31^. Notably, several of the altered species were microbiota-modified bile acids, indicating that the phenotype extends beyond changes in host-derived primary bile acids. These metabolic changes occurred without detectable differences in microbial community composition, supporting a model in which germline mutp53 alters host-microbial bile acid metabolism without major compositional change. DCA, a microbiota-derived secondary bile acid with established pro-inflammatory and tumour-promoting effects, including modulation of inflammatory and immune signalling in cancer^22,23,29^, showed the strongest positive association with distal tumour burden. Moreover, DCA was reduced following microbiota depletion, alongside several other microbiota-modified bile acids.

We next used complementary depletion and transfer approaches to test whether the intestinal microbial environment contributed to distal tumour growth. Microbiota depletion reduced tumour growth in mutp53 mice but not WT mice and was accompanied by reduced intestinal permeability, attenuation of the elevated colonic inflammatory profile, and decreased faecal bile acids. Conversely, transfer of faecal filtrate from mutp53 donors to WT recipients increased tumour growth and recapitulated the colonic inflammatory cytokine profile observed in mutant mice. Together, these experiments demonstrate that the microbiota contributes to the intestinal abnormalities and enhanced distal tumour growth in mutp53 mice, while transferable components of the faecal filtrate are sufficient to reproduce colonic inflammation and enhanced tumour growth in WT recipients.

Our findings complement Kadosh et al. (2020), who showed that a microbiota-derived metabolite can alter mutant-p53 activity within the intestinal epithelium^15^. Our data extend this relationship in the opposite direction, showing that germline mutp53 alters the intestinal environment and its microbiota-associated metabolic output. Thus, these studies suggest bidirectional interactions between mutant p53 and the gut environment and expand the consequences of germline *TP53* mutation beyond tumour-cell-intrinsic defects.

A microbial contribution to cancer risk has also been described in other hereditary cancer syndromes. In familial adenomatous polyposis, colonic biofilms containing colibactin-producing *E. coli* and enterotoxigenic *Bacteroides fragilis* are enriched in patient mucosa and are sufficient to induce inflammation and tumourigenesis when co-colonized in tumour-prone mice ^32^, and colitis-associated tumourigenesis depends on the genotoxic capacity of resident bacteria ^33^. More recent work links *pks* + *E. coli* carriage in FAP polyps to prior colorectal cancer ^34^. Our findings place LFS in this broader context, while differing in that the intestinal phenotype we observe arises without compositional change and affects a tumour outside the gut.

Several limitations should be considered. The subcutaneous MC38 model measures effects on the growth of established transformed cells and therefore does not address spontaneous tumour initiation. Whether the intestinal phenotype influences spontaneous tumour development remains to be determined. In addition, while several bile acids were associated with tumour burden and altered following microbiota depletion, the relative contributions of these and other transferable faecal components to the tumour-promoting phenotype remain to be determined. Finally, 16S sequencing cannot resolve strain-level differences or directly assess microbial function.

These findings raise the possibility that the intestinal environment represents a modifiable component of cancer susceptibility in germline *TP53* carriers. Bile acid metabolism is particularly compelling for further study because multiple species were elevated in the mutant intestine and reduced following microbiota depletion, while the microbiota-derived secondary bile acid DCA showed the strongest positive association with tumour burden. Bile acid profiling in human LFS cohorts will be important to determine whether this metabolic phenotype is conserved in patients and could inform biomarker development or therapeutic targeting. More broadly, our findings suggest that hereditary cancer predisposition genes may influence cancer risk not only through cell-intrinsic defects in tumour suppression, but also by shaping the metabolic and inflammatory state of otherwise non-malignant host tissues.

## Methods

### Mouse models

Wild-type (WT) and heterozygous *Trp53*^R172H/+^ mice on a C57BL/6 background were used in this study. Mice were obtained from Jackson Laboratory (stock number 008183). Both male and female mice were included in all studies. Animals were between 12–20 weeks of age at the start of experiments, depending on the specific experimental protocol. Mice were housed in a facility with 12-hour day and night cycle under specific pathogen-free (SPF) conditions and maintained on a standard 5% irradiated chow diet. Mice were euthanized by carbon dioxide (CO_2_) asphyxiation in accordance with institutional animal care guidelines. All animal procedures were approved by the University Health Network Animal Care Committee and conducted in accordance with approved Animal Use Protocol (AUP) 7039.

### FITC-dextran gut permeability assay

Intestinal barrier integrity was assessed using a fluorescein isothiocyanate (FITC)-dextran 4kDa permeability assay. Mice were fasted from both food and water for 4 hr prior to oral gavage with 200 μL of FITC-dextran (Sigma-Aldrich, cat# 46944-500MG-F) dissolved in sterile phosphate-buffered saline (PBS) at 80 mg/mL, corresponding to a dose of approximately 600 mg/kg. One hour after gavage, blood was collected by terminal cardiac puncture into EDTA tubes (Sarstedt Inc, cat# 41.1395.105). Blood was centrifuged at 1,500 x g for 10 min at 4°C, and the resulting supernatant was further centrifuged at 10,000 x g for 10 min at 4°C to isolate plasma. 50 μL of plasma samples were diluted 1:1 with PBS prior to reading, and resulting concentrations were multiplied by two to back-calculate the undiluted plasma concentration. A standard curve was prepared by serial two-fold dilution of the FITC-dextran gavage stock in PBS (7.8125, 3.90625, 1.953125, 0.976563, 0.488281, and 0.244141 μg/mL). 100 μL of plasma was measured in duplicate using a plate reader (Molecular Devices SpectraMAX Gemini EM) at an excitation wavelength of 485 nm and an emission wavelength of 530 nm. Plasma from untreated, non-gavaged mice was included on each plate as a background control.

### Histology and Immunohistochemistry (IHC)

Large intestinal tissues were harvested, fixed in formalin (Simport Scientific Inc., M961-20FW) for 24 h, and subsequently transferred to 70% ethanol prior to paraffin embedding and staining by STTARR Pathology facility. Paraffin-embedded tissues were sectioned at 5 μm thickness. Haematoxylin and eosin (H&E) staining was performed for histological assessment. Stained sections were imaged using 3D Histech Pannoramic 250 Flash III Dx Digital Scanner at Sickkids Imaging Facility, and quantitative analyses were performed using QuPath software ^35^.

### Flow cytometry

Lamina propria lymphocytes (LPL) were isolated from the large intestine using an established enzymatic digestion protocol ^36^. With some modification of the cited protocol, mice were euthanized and large intestines were removed with two open ends. The colon was rinsed with 3 mL 1 X HBSS (WISENT INC., 311-506-CL, diluted 10 times in Milli-Q water) by using a 18G × 50mm curved gavage needle (Gavageneedle, AFN1850C) and syringe (BD, 309657) to plunge through one end. The large intestine was cut longitudinally using scissors and transferred to 15 mL of 3% media (Table S1) and shaken gently for 10-15s to remove any faecal debris on the intestine. Clean opened tissue was then transferred to 10 mL of strip media (Table S2) and incubated for 20 min at 37°C on a horizontal shaker (New Brunswick Scientific, I24 Incubator shaker) at 80-100 rpm. After the incubation, the tissue was vortexed vigorously for 30s before transferring the tissue to a 70 μm cell strainer (Falcon®, 352350). The flowthrough contains the intraepithelial lymphocytes (IEL). Next, tissue was removed from the strainer and transferred to 15 mL shake media (Table S3) and vortexed vigorously for 30s. The tissue was then transferred to the cell strainer, and flowthrough was collected and combined with the previous flowthrough for all the IEL extracted from the tissue. The remaining tissue was then placed in a 6-well plate (SARSTEDT Inc, 83.3920) with 1X RPMI 1640 (WISENT INC., 350-000-CL) to rinse off residual EDTA from the strip and shake media. After the wash, tissue was cut into 1mm^3^ pieces and incubated in 5 mL digestion media (Table S4, S5) at 37°C for 28 min on a horizontal shaker at 80-100 rpm. After incubation, the tissue was vortexed for 30s and filtered through a cell strainer. The cell strainer was then rinsed with 15 mL of 3% media. This flowthrough contains the LPL required for antibody staining and flow cytometry analysis. LPL suspension was then centrifuged at 450 × *g* for 10 min at 4°C using Eppendorf Centrifuge 5810R. Supernatant was discarded and the pellet was resuspended in 5 mL of 3% media. Resuspended cells were then filtered with a cell strainer and rinsed with 5 mL of 3% media before being centrifuged at 450 × *g* for 10 min at 4°C. The pellet was resuspended with 1-2 mL of FACS buffer (Table S6) and counted with a haematocytometer (Electron Microscopy Sciences, #63510-20) to determine the total cell number. 1 × 10^6^ LPL cells were aliquoted into 5 mL FACS tubes (Falcon®, 352235) and centrifuged at 300 × *g* for 5 min at 4°C. The pellet was resuspended with Zombie Violet^TM^ dye (Table S7, in 1:1250 dilution) as a live/dead discriminator for 15 min at room temperature (RT) in the dark. 1 μL of TruStain FcX (Table S7) was added and incubated for 5 min on ice and in the dark to inhibit the non-specific binding of immunoglobulin to Fc receptor. After that, antibody cocktail (Table S7) was added and incubated for 30 min on ice in the dark. After the incubation, cells were washed with 3 mL of FACS buffer and centrifuged at 300 × *g* for 5 min at 4°C. Samples were then ready for flow cytometry analysis.

Data were acquired on a BD FACSymphony A3 Cell Analyzer, and compensation controls for all fluorochrome-conjugated antibodies were prepared using UltraComp eBeads™ Compensation Beads (Invitrogen™, 01-2222-42), except Zombie Violet™ dye, for which live cells and heat killed cells were used (65°C for 7 min, using Eppendorf ThermoMixer). Analysis of the data was performed on FlowJo software. Cells were gated on singlets and live cells (Zombie Violet^-^), followed by CD45^+^ leukocytes and exclusion of CD19^+^ B cells. Within the CD45^+^CD19^-^ population, myeloid subsets were defined as follows: inflammatory monocytes, Ly6C^hi^F4/80^-^CD11b^+^CD11c^-^; patrolling monocytes, Ly6C^lo^F4/80^-^CD11b^+^CD11c^-^; early inflammatory macrophages, Ly6C^hi^F4/80^+^CD11b^+^CD11c^-^; inflammatory macrophages, Ly6C^hi^F4/80^+^CD11b^+^CD11c^+^; tissue resident macrophages, Ly6C^lo^F4/80^+^CD11b^+^CD11c^+^. These subsets correspond to successive stages of monocyte-to-macrophage maturation in the intestinal lamina propria.

### Immunoblotting

Crypts from the large intestine were isolated with minor alteration of a previously described method.^37^ Large intestine was removed from the euthanized mice, and the remaining faecal matter was pushed gently with forceps on a flat surface. The colon was rinsed with 3 mL 1 X HBSS by using gavage needle and syringe to plunge through one end and cut longitudinally. The intestine was cut into 5 mm pieces and transferred into 25 mL 1X HBSS to incubate for 10 min at RT with gentle shaking during incubation. Tissues were then filtered through 70 μm cell strainer and placed into a fresh 10 mL 1X HBSS with gentle shaking. The wash was done for the second time before transferring the tissue into 10 mL 8 mM EDTA (BioBasic, SD8135) in 1X HBSS and incubated on ice for 30 min. Tissues in EDTA were vortexed for 30s every 10 min during the incubation. After the incubation, tissues were passed through the cell strainer and washed two times with 10 mL 1X HBSS. The eluant was collected and centrifuged at 300 × *g* for 10 min at 4°C. The pellet was collected for downstream immunoblotting.

Pellets containing the large intestine crypts were lysed in 1X RIPA Lysis buffer (Sigma-Aldrich, 20-188) supplemented with protease (Roche, 11836170001) and phosphatase (Roche, 04906845001) inhibitors. 100 μL of lysis buffer was added onto a 20 μL size pellet. Pellets with lysis buffer were incubated on ice for 30 min with occasional vortex during the incubation. After the incubation, the sample was centrifuged at 14,000 × *g* for 10 min at 4°C using Eppendorf Centrifuge 5430R. Supernatant was collected for protein concentration determination using Pierce™ BCA Protein Assay Kit (Thermo Scientific™, A55865) and quantified at 562nm on plate reader (Molecular Devices, VersaMax Plate Reader). Equal amounts of protein (8–20 μg based on the sensitivity of the antibody) were resolved on 4-12% Bis-Tris Plus gels (Invitrogen^TM^, NW04125BOX) in 1X MOPS running buffer (Invitrogen™, B0001) in Invitrogen™ Mini Gel Tank (Invitrogen™, A25977) with PowerPac 3000 Bio-Rad Electrophoresis Power Supply (Bio-Rad, 1655050) set at 160 V for 50 min at RT. PAGE gels were then transferred to PVDF membranes (Cytiva Amersham^TM^ Hybond^TM^, 10600021) in a Bio-Rad Buffer Tank (Bio-Rad, 1658040) submerged in 1X Transfer Buffer (Table S8) at 20 V overnight at 4°C.

The membrane was removed after the transfer and blocked with 5% Non-Fat Powdered Milk (BioBasic, NB0669) in 1X TBST (Table S9) on a rocker (Stovall Life Science Inc., Belly Dancer) for 1 hr at RT. After the incubation, membrane was washed three times 5 min with 1X TBST on the rocker at RT. Membranes were probed with primary antibodies (Table S10) diluted in 5% BSA (WISENT Inc., 800-095-EG) in 1X TBST on a rocker overnight at 4 °C. The next day, membranes were washed three times 5 min with 1X TBST on the rocker at RT to remove any non-binding primary antibody on the membrane. Membranes were then probed with appropriate secondary antibodies (Table S10) on the rocker for 1 hr at RT. Membranes were washed three times 5 min with 1X TBST on the rocker at RT. Protein bands were visualized using ECL™ Prime Western Blotting Detection Reagent (Cytiva Amersham™, 45-010-090) and imaged using the chemiluminescence detection software in ChemiDoc™ MP System (Bio-Rad, 170-8280). Band intensities were quantified using ImageJ software ^38^.

### Intestinal crypt cytokine analysis

Protein lysates extracted from colon crypts described previously were used for cytokine quantification by enzyme-linked immunosorbent assay (ELISA) using commercially available kits from MABTECH and according to the manufacturer’s instructions. Cytokines measured included IL-1β (3317-1H-6), IL-6 (3361-1H-6), TNFα (3511-1H-6), IFNγ (3321-1H-6), IL-10 (3432-1H-6), IL-22 (3376-1H-6), and IL-17a (3521-1H-6). Each sample was analyzed in duplicate and cytokine concentrations were normalized using standard curves provided by the manufacturer. In brief, protein lysates were diluted according to the concentration determined by serial dilutions of the samples to each cytokine. Before the day of the experiment, (1 × 8 Well) Strip Plates (Greiner Bio-One, 762071) were incubated with capture monoclonal antibody (mAB) diluted in 1X PBS (WISENT Inc, 311-010-CL) on a rocker overnight at 4°C. After the coating overnight, the solution was removed and plates were blocked with 0.05% Tween-20 and 0.1% BSA in 1X PBS (incubation buffer) for 1 hr at RT on the rocker. Plates were washed 5 times with 0.05% Tween-20 in 1X PBS (washing buffer). Samples and the standards diluted in incubation buffer were added to the plates and incubated for 2 hr at RT on the rocker and were then washed 5 times with washing buffer. Detection mABs diluted with incubation buffer were added to the plates and incubated on the rocker for 1 hr at RT, then washed 5 times with washing buffer. Streptavidin-HRP diluted in incubation buffer was added and incubated for 1 hr at RT. Washes were done five times with washing buffer before adding TMB substrate (3652-F10) and incubated on a rocker for 15 min at RT and protected from light. After 15 min, 0.1M *H*_2_*SO*_4_ (VWR, BDH7230-1) were added into the substrate to stop the reaction. The optical density of each well was measured with a plate reader (Molecular Devices, VersaMax Plate Reader) at 450 nm and 650 nm. The concentration of each well was calculated using Arigo Bio’s GainData ELISA calculator (www.arigobio.com/elisa-calculator).

### Faecal sample collection

Faecal samples were collected from healthy WT and mutant mice under basal conditions. To minimize environmental contamination and cage-derived variability, mice were individually isolated and transferred to a clean, autoclaved cage for a period of three hours prior to sample collection. Freshly deposited faecal pellets were collected immediately after defecation, snap-frozen, and stored at −80°C until downstream processing. Where indicated, the same faecal samples were used for both microbiome sequencing and untargeted metabolomic analyses, enabling matched comparisons between microbial composition and metabolic output.

### Faecal and plasma metabolomic analysis

Untargeted metabolomic profiling was performed on faecal samples to assess global differences in microbial-derived metabolites between WT and mutant mice. Metabolomic analysis was conducted by The Metabolomics Innovation Centre (TMIC; Calgary, Canada). For systemic metabolite assessment, blood samples were collected as described: mice were warmed under a heat lamp for 5 min prior to blood collection to promote vasodilation. Peripheral blood was collected from mice via saphenous vein bleeding into EDTA-coated tubes (SARSTEDT Inc., 20.1288.100). Samples were centrifuged at 1,500 × *g* for 10 min at 4°C to isolate the plasma. The plasma was then further centrifuged at 10,000 × *g* for 10 min at 4°C to remove residual platelets and cell debris, which was aliquoted and stored at −80°C until analysis. Plasma was shipped on dry ice to TMIC for analysis. Targeted bile acid profiling was performed to quantify individual circulating bile acid species. Sample preparation, metabolite detection, and compound identification were performed by TMIC using established and validated analytical platforms. Resulting metabolite concentrations were used for downstream statistical comparisons between WT and mutant mice.

### DNA extraction from faecal samples

Genomic DNA was extracted from frozen faecal pellets using the AllPrep PowerFecal Pro DNA/RNA Kit (Qiagen, catalogue #80254), with faecal material lysed using vortexing according to the manufacturer’s protocol. DNA yield and purity were assessed using a DeNovix DS-11 series spectrophotometer, and DNA integrity was confirmed by gel electrophoresis (1% agarose, 120V for 45 minutes) prior to submission, as required by the sequencing facility. Extracted DNA was subsequently submitted for 16S rRNA gene amplicon sequencing.

### Microbiome sequencing and bioinformatic analysis

Microbial community composition was assessed by sequencing the V4 hypervariable region of the bacterial 16S rRNA gene. Library preparation and paired-end sequencing were performed by the Centre for Analysis of Genome Evolution & Function (CAGEF), University of Toronto using the Illumina MiSeq platform.

Processed OTU tables and corresponding taxonomic annotations were imported into R statistical software for downstream analysis. All microbiome analyses were conducted using the phyloseq package within the Bioconductor framework ^39^. Alpha diversity was assessed using observed richness, Shannon diversity, and Simpson diversity indices. Beta diversity was evaluated using Bray–Curtis dissimilarity and UniFrac distance metrics and visualized using principal coordinates analysis (PCoA) ^40^.

Statistical comparisons of alpha diversity metrics between WT and mutant mice were performed using the Wilcoxon rank-sum test. Differences in overall microbial community structure were assessed using permutational multivariate analysis of variance (PERMANOVA) ^41^.

### Cell culture

MC38 murine colon adenocarcinoma cells were kindly provided by Dr. Jean Gariépy (University of Toronto, Ontario, Canada). Cells were cultured in DMEM (WISENT Inc., cat # 319-010-CL) supplemented with 10% fetal bovine serum (Cytiva, SH30396.03) and used for tumour challenge between passages 10 and 15. The line was confirmed to be negative for mycoplasma contamination (Charles River Laboratories, UHN Mouse 9 agent CLEAR panels). Cells were cultured at 37°C in a 5% CO_2_ atmosphere (PHCbi, MCO-170AICUVL-PA). Cells were detached using 0.05% trypsin (WISENT Inc., cat # 325-542-EL) for 5 min at 37°C, then washed three times in cold PBS (centrifuged at 300 × g for 5 min), and resuspended at 2.5 × 10^6^ cells/mL. Cell count and viability were confirmed by trypan blue exclusion (WISENT Inc., cat# 609-130-EL) using a cell counter (DeNovix, CellDrop BF) prior to injection.

### Syngeneic tumour model

All animal procedures were approved by the AUP 7039 and performed in accordance with ARC guidelines for the care and use of laboratory animals. Mice were injected subcutaneously in the right flank with 500,000 MC38 cells in 200 μL of PBS using 27G needle (BD, 305109). Animal health and tumour progression were monitored every other day, increasing to daily observations when the tumours approached the experimental endpoint. Tumour volume was measured using calipers (MAXIMUM, #058-1639-0) and estimated using the formula:

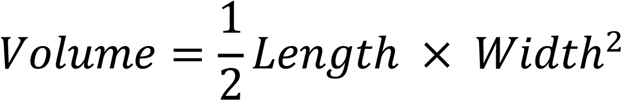

Humane endpoints were strictly enforced when tumour ulceration or necrosis occurred or if the body condition scores fell below an acceptable standard. To maintain cohort consistency, the entire experiment batch was terminated when any tumours exceeded 2000 mm^3^.

### Antibiotic treatment protocol

To deplete the gut microbiota, mice were treated with a broad-spectrum antibiotic cocktail in 0.9% sterile saline (Sigma-Aldrich, S3014-1KG, dissolved with MiliQ water and filtered using 0.2 μm PES membrane (Thermo Scientific^TM^ Nalgene^TM^ Rapid-Flow^TM^, 09741202)), which comprised amoxicillin (1000 mg/L; Cayman Chemical Company, 19188), clavulanic acid (250 mg/L; Toronto Research Chemical, C563751), ciprofloxacin (200 mg/L; Cayman Chemical Company, 14286), vancomycin (500 mg/L; Cayman Chemical Company, 15327), and metronidazole (1000 mg/L; Cayman Chemical Company, 9002409). To ensure consistency across the study, master batch was aliquoted and stored at –20°C, with individual aliquots thawed immediately before each gavage session. The antibiotic cocktail was given four days before the subcutaneous injection of MC38 cells and was maintained once daily until the experimental endpoint. The antibiotic cocktail was administered via oral gavage in 200 μL using a gavage needle. Control group mice received an equivalent volume of sterile saline via oral gavage on the same daily schedule. Following each gavage procedure, animals were monitored for signs of respiratory distress or oesophageal injury to ensure animal welfare.

### Faecal filtrate transplant

Mice were randomly assigned to three experimental groups to investigate the influence of mutant gut microbiome on tumourigenesis. All mice initially received a four-day course of broad-spectrum antibiotic cocktail (as described above) to deplete the endogenous commensal microbiota. Then, mice were treated for an additional three days according to their group assignment:

1. FFT group: received 200 μL of mutant mouse faecal filtrate daily.
2. Antibiotic group: continued daily antibiotic cocktail to maintain depletion.
3. Control group: received 200 μL sterile saline daily as a negative vehicle control.

On the fourth day of the intervention period, all mice were challenged with 500,000 MC38 cells subcutaneously. Daily treatments were maintained until the experimental endpoints.

Faecal filtrate was prepared from donor pellets collected from mutant mice and pooled. Pellets were resuspended in sterile saline at a ratio of 1:1000 (w/v). The suspension was then centrifuged at 4,000 × g for 30 min at 4°C to remove particulate matter. Supernatant was sequentially filtered through 0.45 μm (VWR, 76479-012) and 0.22 μm (Whatman®, WHA69002502) syringe filters to exclude intact bacterial cells.

The sterile filtrate was then aliquoted and stored at −20°C to maintain batch consistency throughout the treatment period. Each treatment dose was thawed immediately before the daily oral gavage. Following each procedure, mice were monitored for clinical signs of distress to ensure animal welfare.

### Statistical analysis

Statistical analyses were performed using GraphPad Prism. Comparisons between two groups were conducted using Welch’s t-test or paired t-test or Wilcoxon ranked-sum test, as appropriate. Data are presented as mean ± SEM unless otherwise indicated. A p value < 0.05 was considered statistically significant, consistent with standard practices in biomedical research. Multiple-comparison corrections using Benjamini-Hochberg False Discovery Rate were applied when appropriate ^42^.

## Acknowledgements

We thank Jodi Lees for laboratory management, Noa Alon for technical assistance, Ann Gong for clinical research project management, and the other members of the Malkin laboratory for helpful discussions. We thank Dr. Anastasia Tikhonova and Dr. John Brumell for their mentorship as members of N.W.Y.O.’s supervisory committee, and for the many helpful discussions that shaped this work. We thank Dr. Jean Gariépy (University of Toronto) for providing the MC38 cell line. We thank Roberto Lopez, Maria Bisa, Vaida Undziene, Andrea Archila, Jason Alverio and Audrey Pang of the Animal Resource Centre (University Health Network) for animal husbandry and veterinary support. We thank the STTARR Innovation Centre (University Health Network) for histology processing and imaging support, the SickKids Imaging Facility for slide scanning, the Centre for the Analysis of Genome Evolution and Function (University of Toronto) for 16S rRNA gene sequencing, and The Metabolomics Innovation Centre (Calgary, Canada) for metabolomic profiling.

## Author contributions

Conceptualization: N.W.Y.O., D.M.

Methodology: N.W.Y.O., N.W.F.

Investigation: N.W.Y.O., N.W.F., C.G., P.R.Q., Y.P., R.T.

Formal analysis: N.W.Y.O.

Visualization: N.W.Y.O.

Writing – original draft: N.W.Y.O.

Writing – review & editing: N.W.Y.O., N.W.F., D.M.

Supervision: D.M.

Funding acquisition: D.M.

## Funding

This work was supported in part by a New Frontiers Program Project Grant (#1081) of the Terry Fox Research Institute and a Foundation Scheme Grant From the Canadian Institutes for Health Research (to DM). DM is supported in part by the CIBC Children’s Foundation Chair in Child Health Research.

## Competing interests

The authors declare no competing interests.

## Notes

### Competing Interest Statement

The authors have declared no competing interest.

### Summary of Updates

Changes have been made on the author list, and statistic update on Figure 1.

