## Supplementary Figure for "A microbiota-dependent gut bile acid axis links germline mutant p53 to distal tumour growth"

Table S1: Composition of 3% media, used for tissue rinsing, faecal debris removal, and resuspension of lamina propria lymphocytes.

| Reagents | Company | Catalog # | Stock conc | Final conc | Volume |
|---|---|---|---|---|---|
| RPMI 1640 | WISENT | 350-000-CL | | | 470 mL |
| Pen/Strep | WISENT | 450-201-EL | 10,000 U/mL | 100 U/mL | 5 mL |
| FBS | Cytiva | | 100% | 3% | 15 mL |
| HEPES | WISENT | 330-050-EL | 1M | 20 mM | 10 mL |

Table S2: Composition of strip media, used for intraepithelial lymphocyte removal from intestinal tissue.

| Reagents | Company | Catalog # | Stock conc | Final conc | Volume |
|---|---|---|---|---|---|
| 3% media | | | | | 9.89 mL |
| EDTA | Bio Basic | SD8135 | 0.5 M | 5 mM | 100 µL |
| DTT | Bio Basic | DB0058 | 1 M | 1 mM | 10 µL |

Table S3: Composition of shake media, used for intraepithelial lymphocyte removal from intestinal tissue.

| Reagents | Company | Catalog # | Stock conc | Final conc | Volume |
|---|---|---|---|---|---|
| RPMI 1640 | WISENT | 350-000-CL | | | 483 mL |
| Pen/Strep | WISENT | 450-201-EL | 10,000 U/mL | 100 U/mL | 5 mL |
| EDTA | Bio Basic | SD8135 | 0.5 M | 2 mM | 2 mL |
| HEPES | WISENT | 330-050-EL | 1 M | 20 mM | 10 mL |

Table S4: Composition of 0% media, used as the base for digestion media preparation.

| Reagents | Company | Catalog # | Stock conc | Final conc | Volume |
|---|---|---|---|---|---|
| RPMI 1640 | WISENT | 350-000-CL | | | 479.75 mL |
| Pen/Strep | WISENT | 450-201-EL | 10,000 U/mL | 100 U/mL | 5 mL |
| B-mercapto-ethanol | Sigma-Aldrich | M6250-100ML | 100 mM | 50 µM | 250 µL |
| HEPES | WISENT | 330-050-EL | 1 M | 20 mM | 10 mL |
| Sodium pyruvate | WISENT | 600-110-EL | 100 mM | 1 mM | 5 mL |

Table S5: Composition of digestion media, used for enzymatic digestion of intestinal tissue to release lamina propria lymphocytes.

| **Reagents** | **Company** | **Catalog #** | **Stock conc** | **Final conc** | **Volume** |
|---|---|---|---|---|---|
| 0% Media | | | | | 5.85 mL |
| Liberase TL | Roche | 05401119001 | 5 mg/mL | 100 μg/mL | 120 μL |
| DNase I | Roche | 10104159001 | 100 mg/mL | 500 μg/mL | 30 μL |

Table S6: Phosphate-buffered saline (PBS) supplemented with 2% fetal bovine serum (FBS).

| **Reagents** | **Company** | **Catalog #** | **Stock conc** | **Final conc** | **Volume** |
|---|---|---|---|---|---|
| 1X PBS | WISENT | 311-010-CL | | | 490 mL |
| FBS | Cytiva | SH30396.03 | 100% | 2% | 10 mL |

Table S7: Flow cytometry antibody panel used for lamina propria lymphocyte staining.

| **No.** | **Antibody** | **Fluorescent dye** | **Company** | **Catalog #** | **Volume per sample** | **Target** |
|---|---|---|---|---|---|---|
| 1 | CD45 | APC/Cy7 | Biolegend | 103116 | Diluted in 1:8 and used 1 μL | All immune cells |
| 2 | NK1.1 | R718 | BD | 567380 | 2.5 μL | NK cells |
| 3 | CD11b | APC | Biolegend | 101211 | 1 μL | Myeloid |
| 4 | Ly-6C | PE | Biolegend | 128007 | 1 μL | Myeloid - Particularly macrophages |
| 5 | TCR-b | PerCP/Cy5.5 | Biolegend | 109227 | 1.25 μL | T-cell (Helper and cytotoxic) |
| 6 | CD11c | KIRAVIA blue | Biolegend | 117363 | 1 μL | Myeloid - Particularly macrophages |
| 7 | TCRyd | BV605 | Biolegend | 118129 | 1 μL | T-cell (tissue resident) |
| 8 | CD19 | BUV395 | BD | 563557 | 3 μL | B cells |
| 9 | F4/80 | BUV737 | BD | 749283 | 2 μL | Myeloid - Particularly macrophages |
| 10 | Zombie violet | BV421 | Biolegend | 423114 | Diluted in 1:1250 (100-total Abs-TruStain Fc/X) μL | Live/dead |
| 11 | TruStain FcX | | Biolegend | 101320 | 1 μL | Prevent non-specific antibody binding |

Table S8: Composition of 1X Transfer Buffer

| Reagents | Company | Catalog # | Stock conc | Final conc | Volume |
|---|---|---|---|---|---|
| Tris-Glycine (TG) Buffer | WISENT | 880-560-LL | 10X | 1X | 100 mL |
| MiliQ water | | | | | 700 mL |
| Methanol | Thermo Fischer Scientific | A4124 | 100% | 20% | 200 mL |

Table S9: Composition of 1X Tris-buffered saline Tween-20 (TBST)

| Reagents | Company | Catalog # | Stock conc | Final conc | Volume |
|---|---|---|---|---|---|
| TBS | Bio Basic | UA0045 | 20X | 1X | 50 mL |
| MiliQ water | | | | | 949 mL |
| Tween-20 | Bio Basic | 9005-64-5 | 100% | 0.1% | 1 mL |

Table S10: Primary antibodies used for western blotting

| Reagents | Host species | Company | Catalog # | Dilution | Diluent |
|---|---|---|---|---|---|
| p65 | Rabbit | Cell Signaling Technology (CST) | 8242S | 1:1000 | 5% BSA in 1X TBST |
| p-p65 | Rabbit | CST | 3033S | 1:500 | 5% BSA in 1X TBST |
| β-actin | Rabbit | CST | 4967S | 1:2000 | 5% BSA in 1X TBST |
| Anti-rabbit IgG | Goat | CST | 7074S | 1:2000 | 5% BSA in 1X TBST |

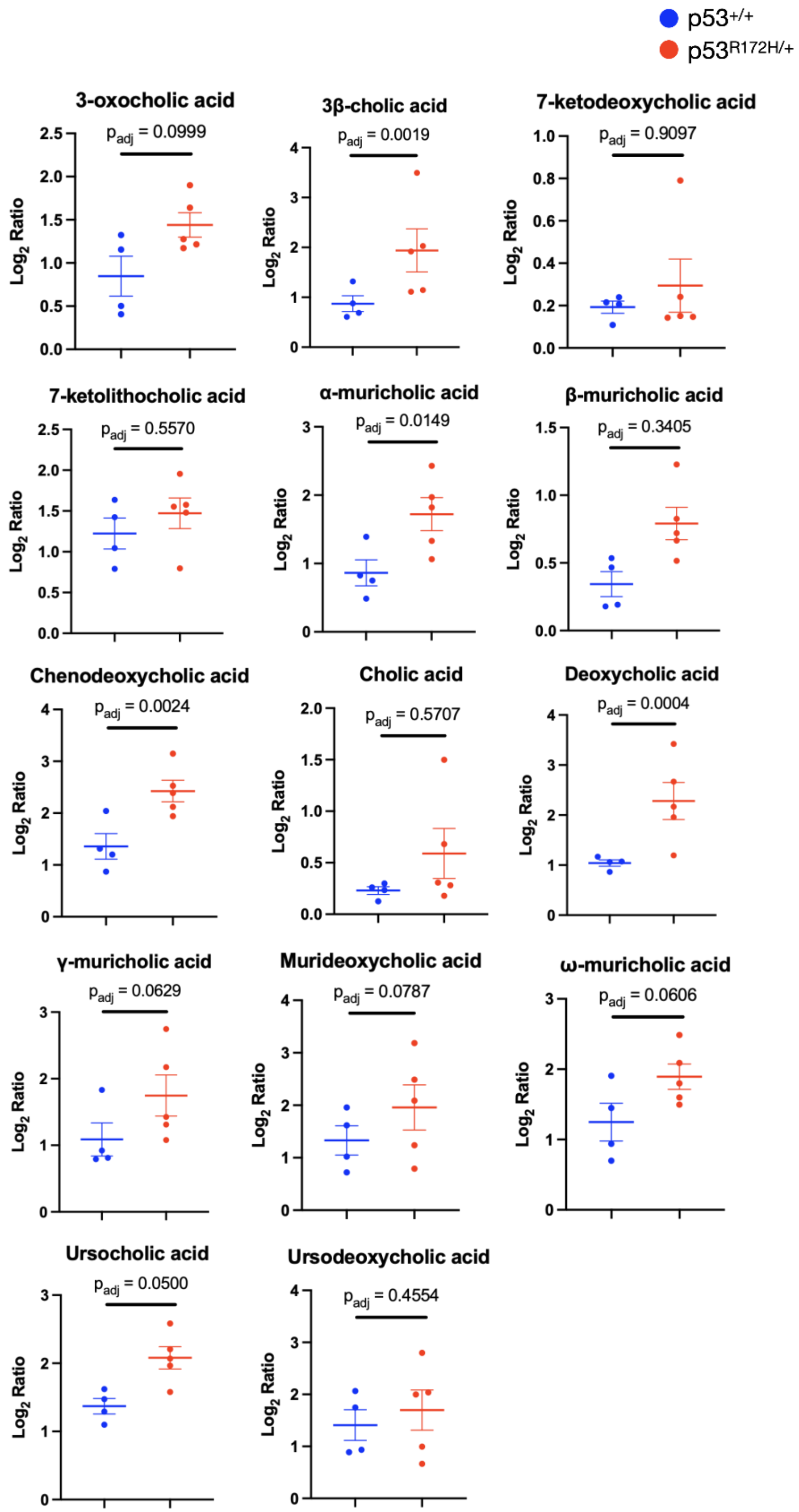


**Supplementary Fig. S1: Individuals sample-level faecal bile acid abundance by species.** $Log_2$ ratio of relative peak intensity (mutant relative to WT) for each of the 14 bile acid species identified within the untargeted faecal metabolomic dataset (see Figure 4.2). Points show individual mice (WT, blue n = 4; mutant, red, n = 5); horizontal bars show mean ± SEM. Adjusted p-values (BH-FDR) are shown above each panel.

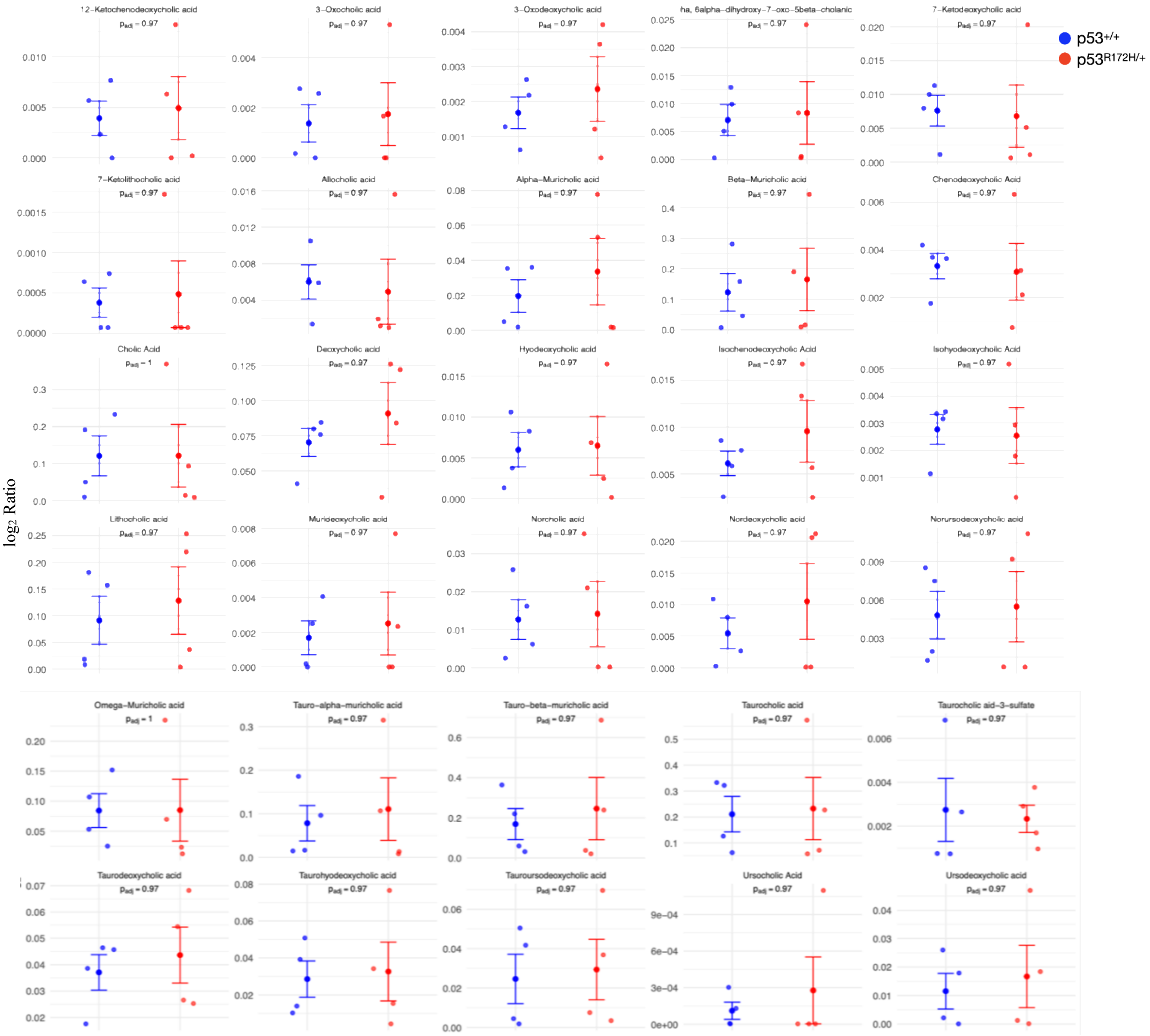


**Supplementary Fig. S2: Individual sample-level plasma bile acid abundance by species.** $Log_2$ ratio (mutant (red, n = 4) relative to WT (blue, n = 4)) for each of the 30 targeted plasma bile acid species shown in Figure 4.4. Points show individual mice; larger points and error bars show mean ± SEM per group. Adjusted p-value (Benjamini-Hochberg correction, Welch's t-test per species) are shown above each panel; $p_{adj} \geq 0.97$ for all species.

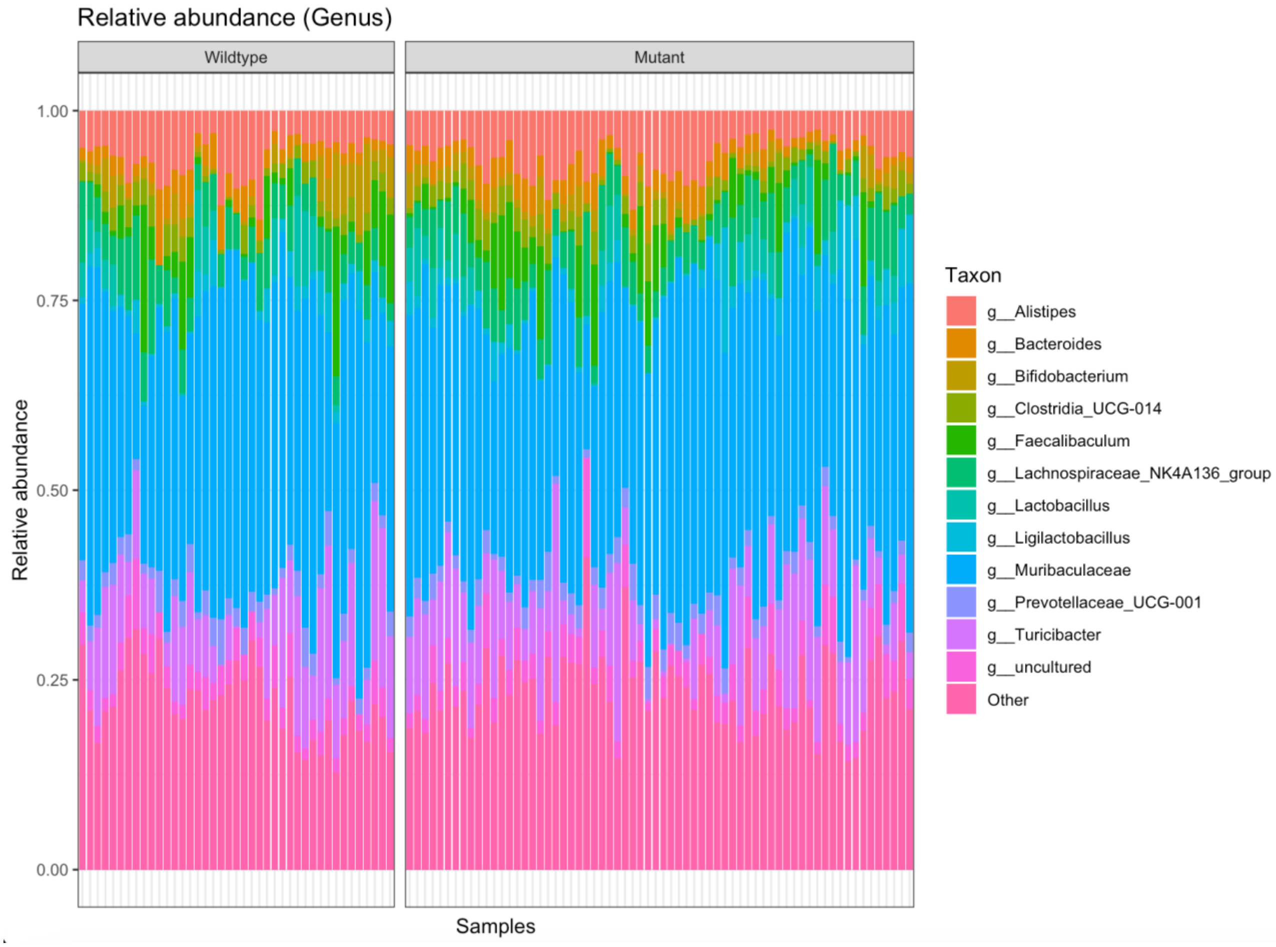


**Supplementary Fig. S3: Genus-level taxonomic composition does not differ by genotype.** Relative abundance of the 13 most abundant bacterial genera in WT (n = 41) and mutant (n = 61) faecal samples. Each bar represents one sample; samples are grouped by genotype. “Other” includes genera below threshold (1% relative abundance).

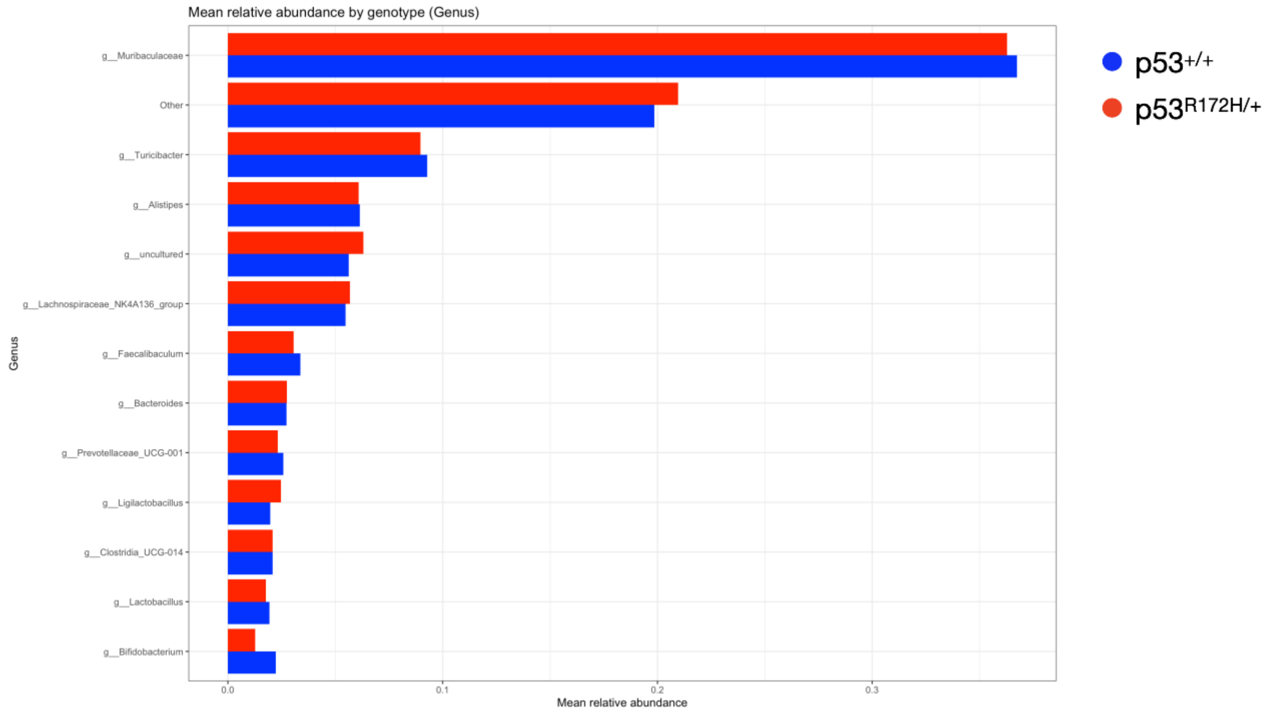


**Supplementary Fig. S4: Mean genus-level relative abundance by genotype.** Mean relative abundance (±SEM) of each genus in WT (blue, n = 41) and mutant (red, n = 61) samples, ranked by overall abundance. No genus differed significantly between genotype (Wilcoxon ranked-sum test, BH-FDR-correct q >X).

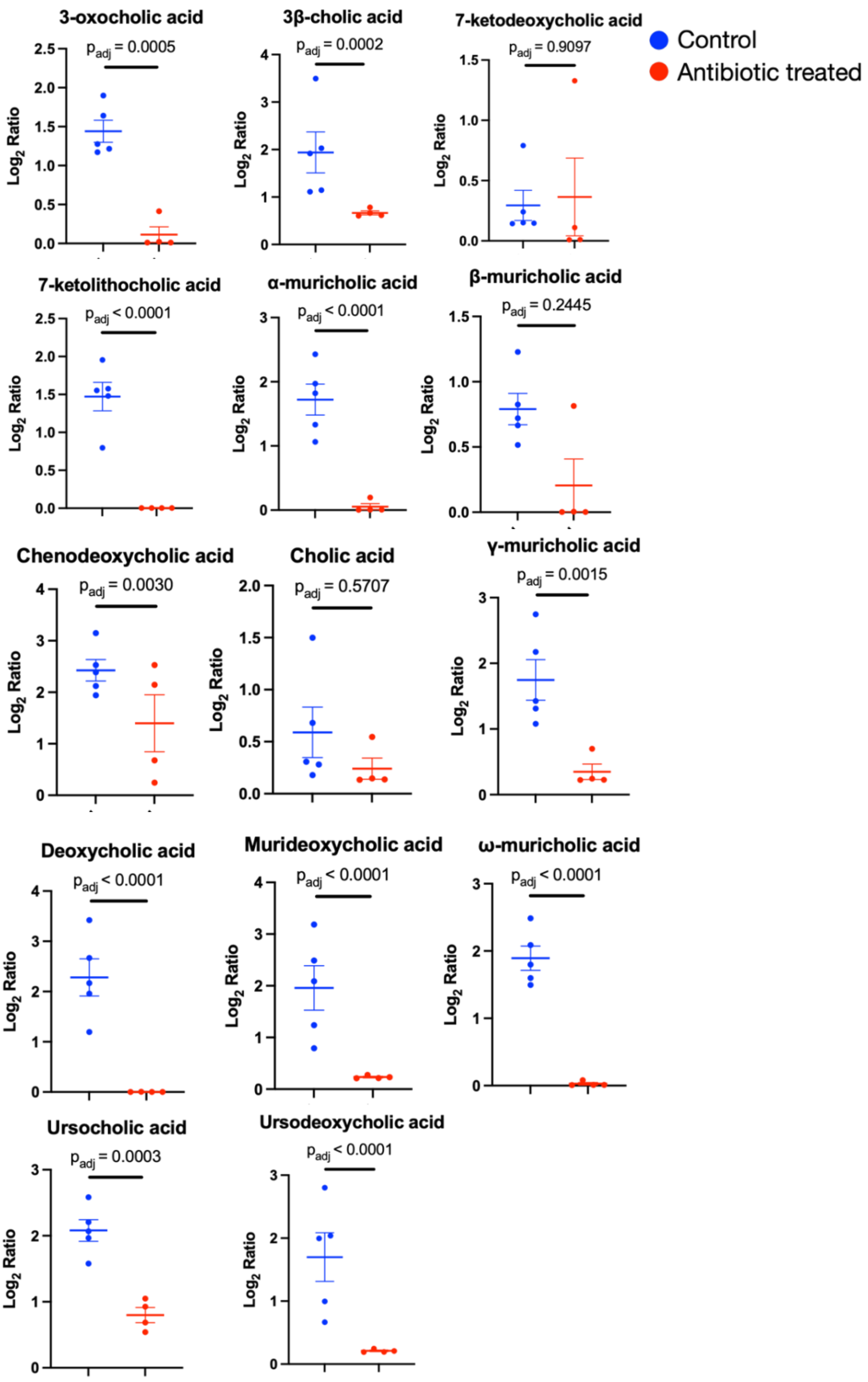

Control
Antibiotic treated
3-oxocholic acid
p_adj = 0.0005
3β-cholic acid
p_adj = 0.0002
7-ketodeoxycholic acid
p_adj = 0.9097
7-ketolithocholic acid
p_adj < 0.0001
α-muricholic acid
p_adj < 0.0001
β-muricholic acid
p_adj = 0.2445
Chenodeoxycholic acid
p_adj = 0.0030
Cholic acid
p_adj = 0.5707
γ-muricholic acid
p_adj = 0.0015
Deoxycholic acid
p_adj < 0.0001
Murideoxycholic acid
p_adj < 0.0001
ω-muricholic acid
p_adj < 0.0001
Ursocholic acid
p_adj = 0.0003
Ursodeoxycholic acid
p_adj < 0.0001
Log2 Ratio

**Supplementary Fig. S5: Individual faecal bile acid comparisons underlying Fig. 4G.** Faecal $\log_2$ ratio of 14 bile acid species in mutant mice, control (blue) vs. antibiotic-treated (red) ($n = 5$ per group). Statistical significance was assessed by one-way ANOVA with BH correction for multiple comparisons; adjusted p-values are shown for each species. Data are mean ± SEM.

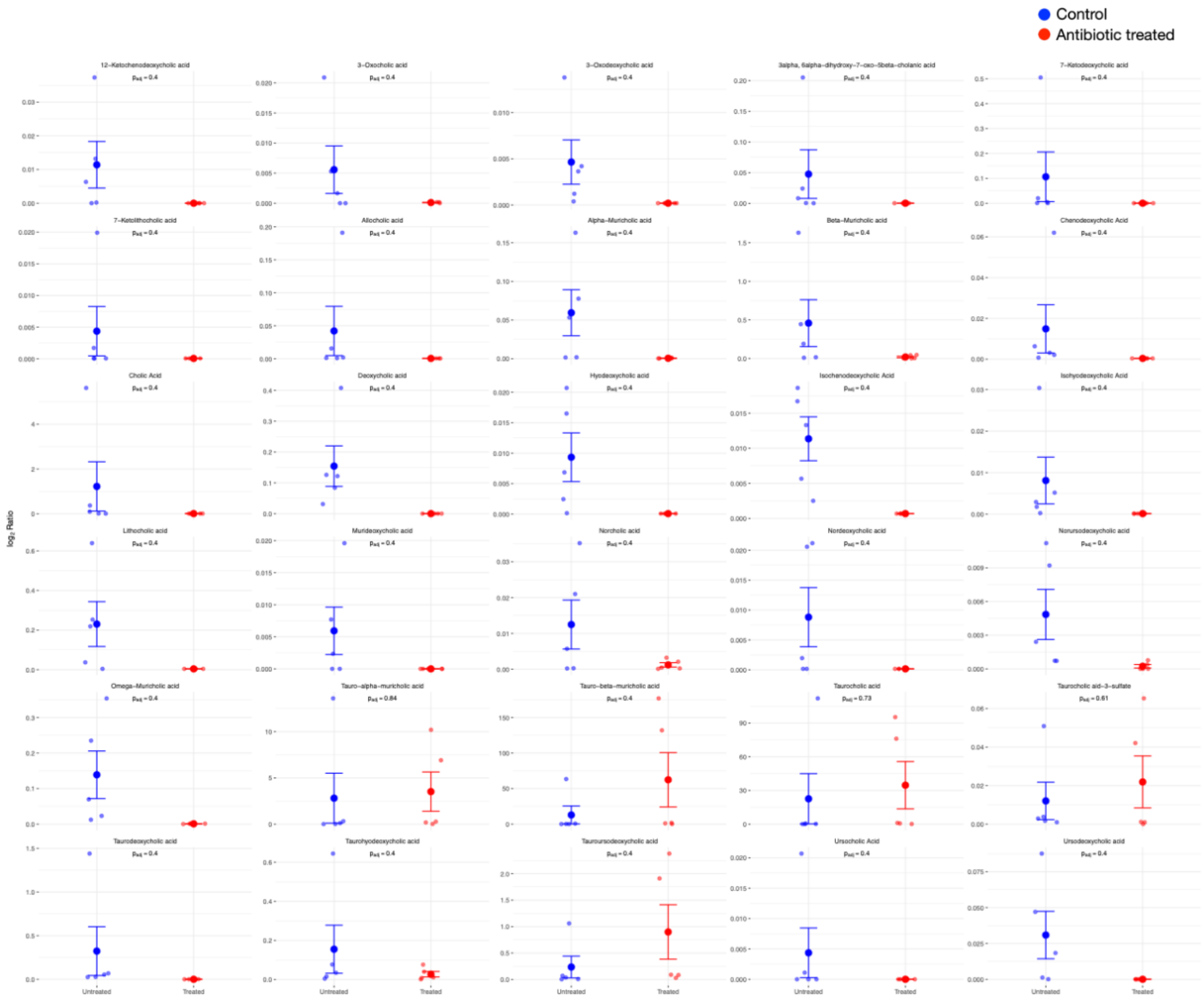


**Supplementary Fig. S6: Individual plasma bile acid comparisons underlying Fig. 4H.** Plasma $\log_2$ ratio of 30 targeted bile acid species in mutant mice, untreated control (blue) vs. antibiotic-treated (red) (n = 5 per group). Statistical significance was assessed by one-way ANOVA with BH correction for multiple comparisons across all 30 species; adjusted p-values are shown for each species, ordered by ascending raw p-value. No species reached significance at $p_{adj} < 0.05$. Data are mean ± SEM.
